# A practical PPE decontamination method using warm air and ambient humidity

**DOI:** 10.1101/2020.11.12.380196

**Authors:** Jesse J. Kwiek, Christopher R. Pickett, Chloe A. Flanigan, Marcia V. Lee, Linda J. Saif, Jeff Jahnes, Greg Blonder

## Abstract

Despite a year-and-a-half of Sars-CoV-2 pandemic experience, personal protective equipment (PPE) remains in short supply^1^. Current decontamination methods are complex, slow, expensive and particularly ill-suited for low to middle income nations where the need is greatest. We propose a low temperature, ambient humidity decontamination method (WASP-D) based on the thirty minute or less half-life of Sars-CoV-2 (and other common pathogens) at temperatures above 45°C, combined with the observation that most PPE are designed to be safely transported and stored at temperatures below 50°C. Decontamination at 12 hours, 46°C (115°F) and ambient humidity should consistently reduce SARS-CoV-2 viral load by a factor of 10^6^, without negatively affecting PPE materials or performance.

## Introduction

In the midst of a pandemic, Personal Protective Equipment (PPE) are a critical line of first defense. Prior to the development of vaccines and symptom treatments, PPE limits the spread of disease and protects the lives of vital healthcare workers. Unfortunately, PPE manufacturing capacity and stockpiles persistently lag demand^2^, resulting in needless deaths and severe economic disruption. Despite best efforts to expand capacity, this PPE access gap continues in developed economies with strong healthcare systems and is exacerbated in low to middle income nations (LMIN)^3^.

One potential stop-gap measure is to safely and effectively decontaminate disposable PPE for re-use. A small cohort of methods have been evaluated and approved for use by the CDC and WHO for N95 masks (knowledge here is rapidly evolving-see N95Decon.org or the CDC^4^ for current information). These include UV-C irradiation, vaporized hydrogen peroxide, dry and moist heat^5^.

While viable, these methods fail to work uniformly on all brands of PPE^6^, require costly electronics/pumps /renewables and maintenance, or do not scale to high capacity. For example, UV-C is a broad-spectrum antimicrobial, but cannot penetrate into the shadows of folded PPE which may harbor viable pathogens. Some UV systems produce ozone which breaks down^7^ plastic surfaces. Chemical treatment such as peroxide vapor is both toxic and requires specialized equipment and staff to operate at large scale. To avoid inadvertent high-temperature thermal degradation from exposed 1500°F electric-heating elements, moist heat (60% RH and 165°F/74°C) ovens must be specially designed to block infra-red radiation^8^. High heat also softens elastomeric straps, which may not return to their original taut lengths, or may reduce the material’s ultimate tear strength.

N95 masks, in particular, contain an inner layer that is electrostatically charged to enhance particle collection efficiency without impacting breathability. This layer is easily neutralized by many wet sanitizers, such as soap and water or autoclaving, making re-use impossible. Other PPE, such as gowns or face shields, can be decontaminated in liquid disinfectants, though may be damaged by physical impact and the process itself is highly labor intensive and residual moisture may encourage microbial growth.

### Our goal is to devise and test an accessible decontamination protocol suited for LMIN^9^

We are seeking a decontamination, not a sterilization, methodology. The key metric is reducing viral burden and associated clinic or hospital pathogens to manageable levels. We recognize that some pathogens, e.g. mesophiles such as Listeria^10^ and methicillin-resistant Staphylococcus aureus (MRSA)^11^, have an active growth rate upper temperature bound that just extends into the WASP-D range. And that no lab test, including this study and associated literature studies, can eliminate the possibility of post-WASP-D re-contamination. But we also note the world is filled with pathogens-every breath^12^ we take inhales e-coli, clostridium botulinum and various actinobacteria, along with air pollution and other toxic fumes.

### Reduction from harm, not elimination of all possible risks, is the goal in a pandemic

Such a protocol would ideally meet the following criteria:

1. Achieve at least 10^6^ reduction in common disease pathogens.
2. Inexpensive.
3. Work on all brands and all types of PPE without damage.
4. Tolerant of intermittent power outages.
5. Constructed locally.
6. Minimize training to operate.
7. Fail-soft.

Simply “waiting” at room temperature has been demonstrated to reduce SARS-CoV-2 by three orders of magnitude over seven days^13^. However, according to the CDC, six orders of magnitude reduction is typically required for safe re-use^14^. Characteristically, pathogens exhibit a thermally activated degradation rate, but the literature contains few reliable measurements of warm temperature viral inhibition. Pathogen half-life depends on a number of external factors, including temperature, humidity, exposure to light (particularly UV), encasing media (e.g. saliva, mucous, water, …) and substrate^15^ (metal, plastic, mesh, air..). While Sars-CoV-2 research continues to advance, literature results to-date indicate a factor of four or more variation^16^ in half-life depending on these external factors. Thus, any practical system must err on the side of caution regarding temperature and humidity.

*The purpose of our study is to determine if low-temperature warming of viruses can achieve a six-order of magnitude viral load reduction. Here we test the ability of low-temperature warming to inactivate two surrogate viruses – bovine coronavirus, a beta coronavirus, like SARS-CoV-2 (enveloped) that infects cattle, and MS-2, a small, non-enveloped virus that infects E*.*coli*.

By far the simplest solution would be a combination of time and warm heat at ambient humidity-**W**arm **A**nd **S**low **P**athogen-**D**econtamination, or WASP-D(T) where “T” is the decontamination temperature. In WASP-D, PPE are placed in a large closed room or container. Warm heat (43°C-50°C=110°F-122°F, nominally 46°C as a safety factor) and ambient air ventilation are introduced for 12 hours, following a protocol suggested in Appendix II. After 12 hours, the amount of infectious virus is reduced by a factor of 10^6^. The same protocol suppresses many molds and fungi which lead to unpleasant mask odors.

Effective, universal, low-tech, scalable and no consumables.

#### WASP-D Parameter Choices

There are three key parameters in this study. First, the choice of warm heat at 43°C-50°C. Second, as moisture affects pathogen viability, a relative-humidity characteristic of affected communities. And third, a pathogen reduction of 10^6^ or greater. Decontamination cycle time is a dependent variable necessary to achieve the above objectives.

#### Warm temperatures

Most PPE are manufactured from plastic. For example, melt spun polypropylene for the N95 electrostatic filter or high-density polyethylene in a Tyvek body suit. Cellulosic padding, natural or thermoplastic elastomers and small metal snaps may be part of the design. In addition, proprietary hydrophilic coatings to repel blood or bodily fluids, anti-microbials and a host of unknown and transient features may be present. Each material will experience its own specific thermally activated failure mechanism and degradation pathways.

Testing every brand of PPE is impractical. However, almost all PPE are intended to be shipped from manufacturer to end-user. Like most products, they are designed to withstand standard shipping and storage conditions. Experimental studies^17^ of containerized shipping containers and trucks indicate 50°C is a typical upper bound (with rare excursions to 60°C. Note at higher temperatures, such as 70°C, N95 masks degrade^18^). Consistent with these findings, many product companies (including 3M for N95 masks^19^) recommend 50°C as the maximum safe storage and shipping temperature. While conservative (some PPE are more accepting of higher temperatures, and future decontamination-grade PPE may be more heat tolerant), we chose 50°C or below as likely to preserve PPE efficacy across all commercial products, irrespective of designs.

#### Humidity

To avoid the cost and complexity of humidity-controlled ovens, the WASP-D protocol draws in ambient air to gently ventilate the heated storage/decontamination chamber. Since the chamber is hotter than ambient, and hot air can “hold” higher levels of moisture and the relative humidity declines. Ventilation also dries out sweat, limiting mold and fungal growth. But until dried out, evaporative cooling lowers^20^ PPE surface temperature (see Appendix I), so the protocol compensates by extending the decontamination cycle.

Climate varies widely across the globe-thus we considered three limiting cases: Boston, Rio de Janeiro and Lagos. By holding the partial pressure of ambient humidity constant as it warms to ventilate the chamber, we find:

| Location | Median temperature | Relative humidity of ambient inlet air at median temperature | Relative humidity at 43°C during cycle |
| --- | --- | --- | --- |
| Boston | 70°F (21°C) | 60% | 17% |
| Rio de Janeiro | 81°F (27°C) | 80% | 22% |
| Lagos | 90°F (32°C) | 80% | 44% |

For our initial experiments, we chose 22%RH (@43°C) as a representative average.

There is some evidence^21^ ambient humidity levels under certain conditions are protective to the virus, especially when the virus is contained in a saliva droplet. Again, WASP-D incorporates additional time as a “guard-band” in compensation.

#### Pathogen Reduction

For safety and expediency, bovine coronavirus, a beta coronavirus like SARS-CoV-2, and MS-2 were chosen as surrogate viruses. As the EPA notes, “According to this hierarchy, if an antimicrobial product can kill a small, non-enveloped virus it should be able to kill any large, non-enveloped virus or any enveloped virus. Similarly, a product that can kill a large, non-enveloped virus should be able to kill any enveloped virus.”^22^

## Experimental Method

### Material testing

N-95 test coupons measuring approximately 1cm^2^ were placed into a single well of a 12-well dish, in triplicate. Over the course of two experiments, an average of 1.2 × 10^6^ TCID_50_ units of BCoV-mebus in minimum essential medium (MEM) containing artificial saliva^23,24^ was spotted onto each test coupon. Each 12-well dish were placed in their respective temperature incubators for the desired time points. Following incubation, N-95 test coupons were incubated with one mL of medium and rocked at room temperature for ten minutes to recover infectious virus.

### Bovine Coronavirus infectivity assay

Madin Darby Bovine Kidney (MDBK) cells were maintained in advanced minimal essential medium (AMEM, Gibco) supplemented with 5% heat-inactivated Fetal Bovine Serum (FBS), 2 mM L-Glutamine (Gibco), and 1% Antibiotic/Actinomycotic cocktail (Gibco) ^25^. The BCoV-Mebus (GenBank: U00735.2) strain was used^26^. Median tissue culture infectious dose (TCID_50_) assays were performed according to published protocols. To detect cytopathogenicity (CPE) caused by the virus, BCoV-infected and uninfected (control) MDBK cells were imaged in a SpectraMax Imaging Cytometer (Molecular Devices) at a 5-millisecond exposure. Negative controls, which included uninfected MDBK cells and MDBK cells incubated with MEM exposed to a N-95 test coupon, were negative for CPE induction. TCID_50_ values were calculated using the Reed-Muench method^27^.

### MS-2 plaque assay

Plaque assays were performed using MS-2, a non-enveloped bacteriophage, to test its ability to survive and infect *E. coli* at the selected temperatures and time durations (*Escherichia coli* bacteriophage MS2 ATCC® 15597B1™). A saliva and MS-2 mixture was spotted onto each test coupon. MS-2 was recovered and eight, 10-fold dilutions were made. Ten microliters of each dilution were spotted, and an agar overlay was added^24^. After a 24-hour incubation period, plaques were counted, and the titer of MS-2 was calculated.

## Results

After 12 hours at 43°C both MS-2 and BoCoV viral load are projected to decline by a factor of 10^6^ (Figure 1). At room temperature (22°C) levels of infectious MS-2 and Bovine Coronavirus decline appears to slow. These results are in rough agreement with a variety of lower temperature measurements made by other groups under different conditions^16^. A viral load reduction of 10^6^ is equivalent to 20 half-lives, so if the half-life at 43°C is around a half-hour, ten hours (20*0.5) will be sufficient to reduce by 10^6^. While not definitive, given the approximately exponential decrease in life-time with temperature by about a factor of 3 per decade Celsius^25^, the WASP-D protocol recommends 46°C and 12 hours.

**Figure 1.**
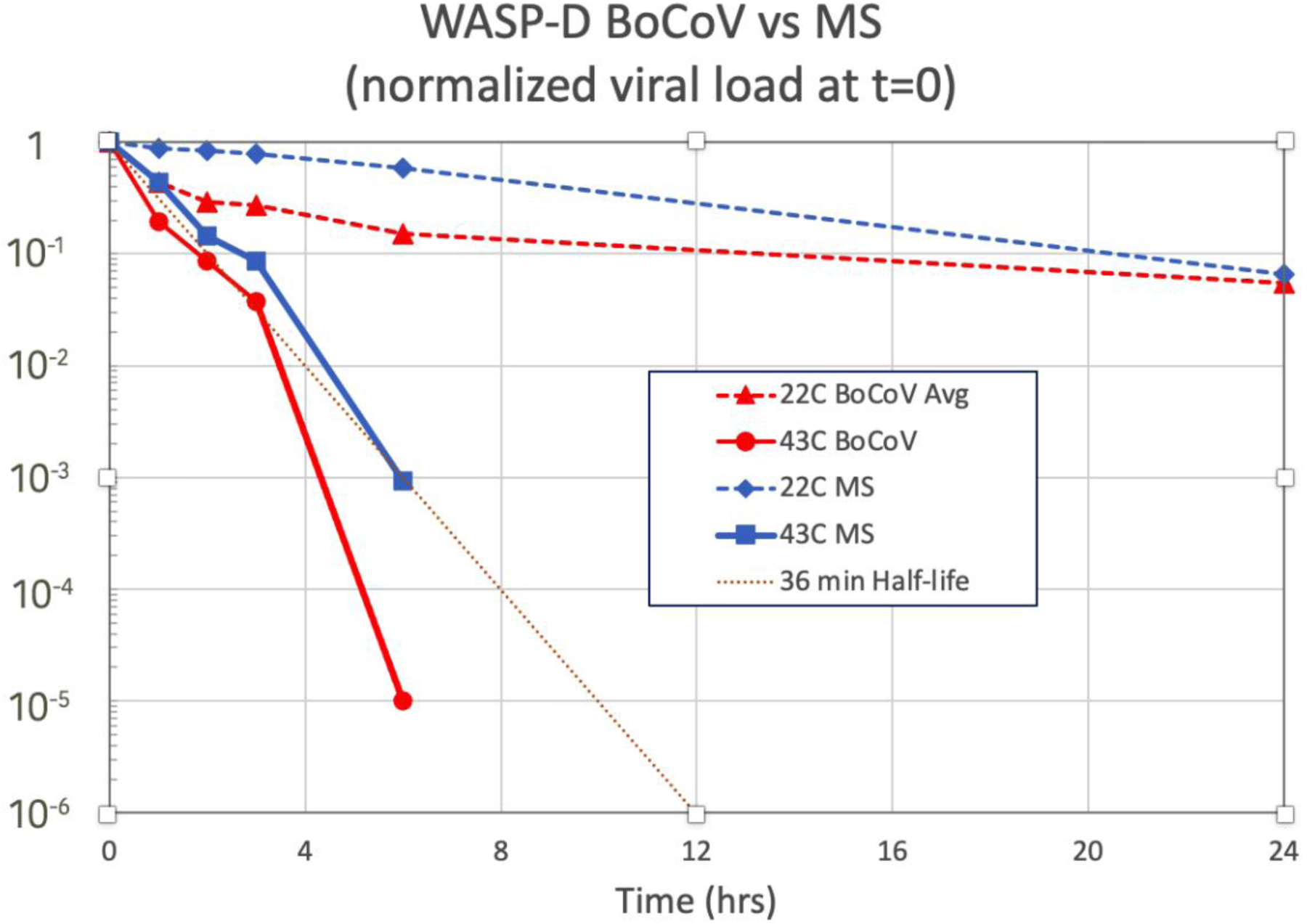
The 22°C Bovine Coronavirus (BoCoV) curve is an average of two experiments. MS = MS-2 bacteriophage. The dotted line denotes an exponential 12 hour reduction by 10^6^.

**Figure 2.**
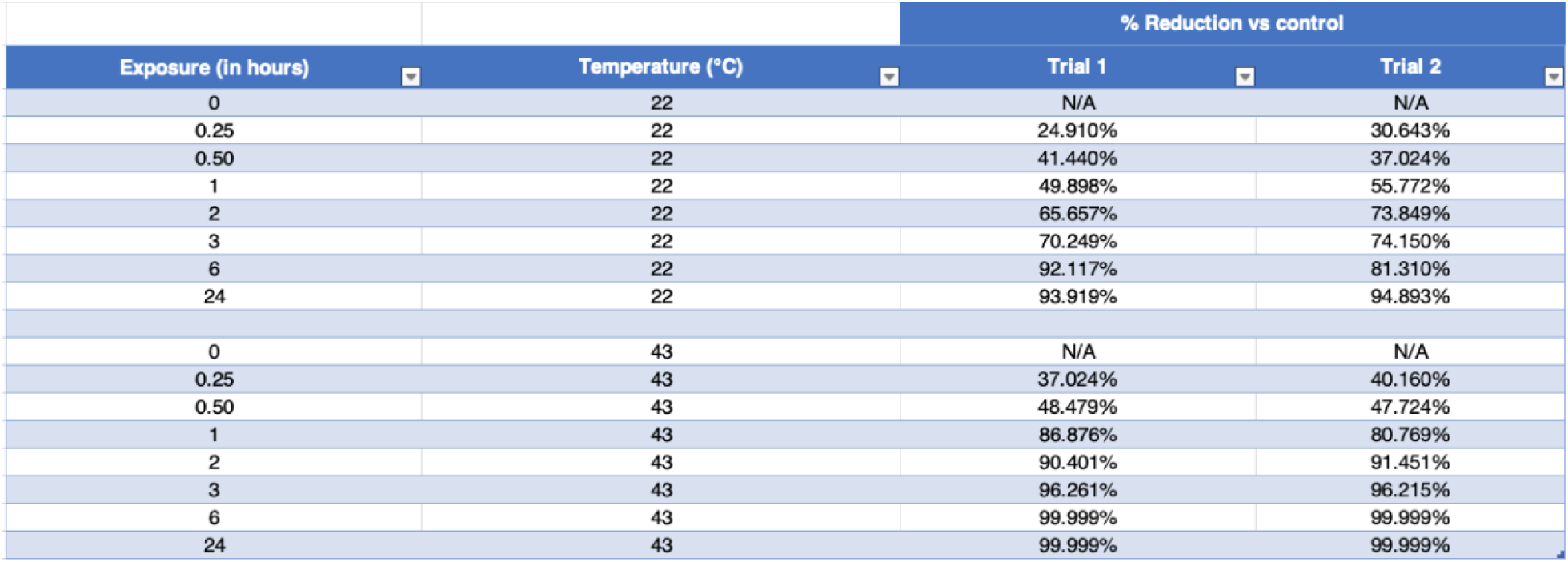
Heat reduces Bovine Coronavirus infectivity. Bovine Coronavirus (strain Mebus) was heated at 22°C or 43°C for up to twenty-four hours. Following heat treatment, Bovine Coronvirus was incubated with MDBK cells for 48-hours, cyotopathogenecity (CPE) was scored, and median tissue culture infectivity (TCID_50_) was calculated. The TCID_50_ assay has a limit of detection of 10 TCID_50_ units (here defined as CPE detected in all four replicates. If we define limit of detection (LoD) as any of the replicates showing CPE, then the LoD is 1 TCID_50_ units. No infectious virus was detected following exposure to 43°C heat for six or 24 hours.

**Figure 3.**
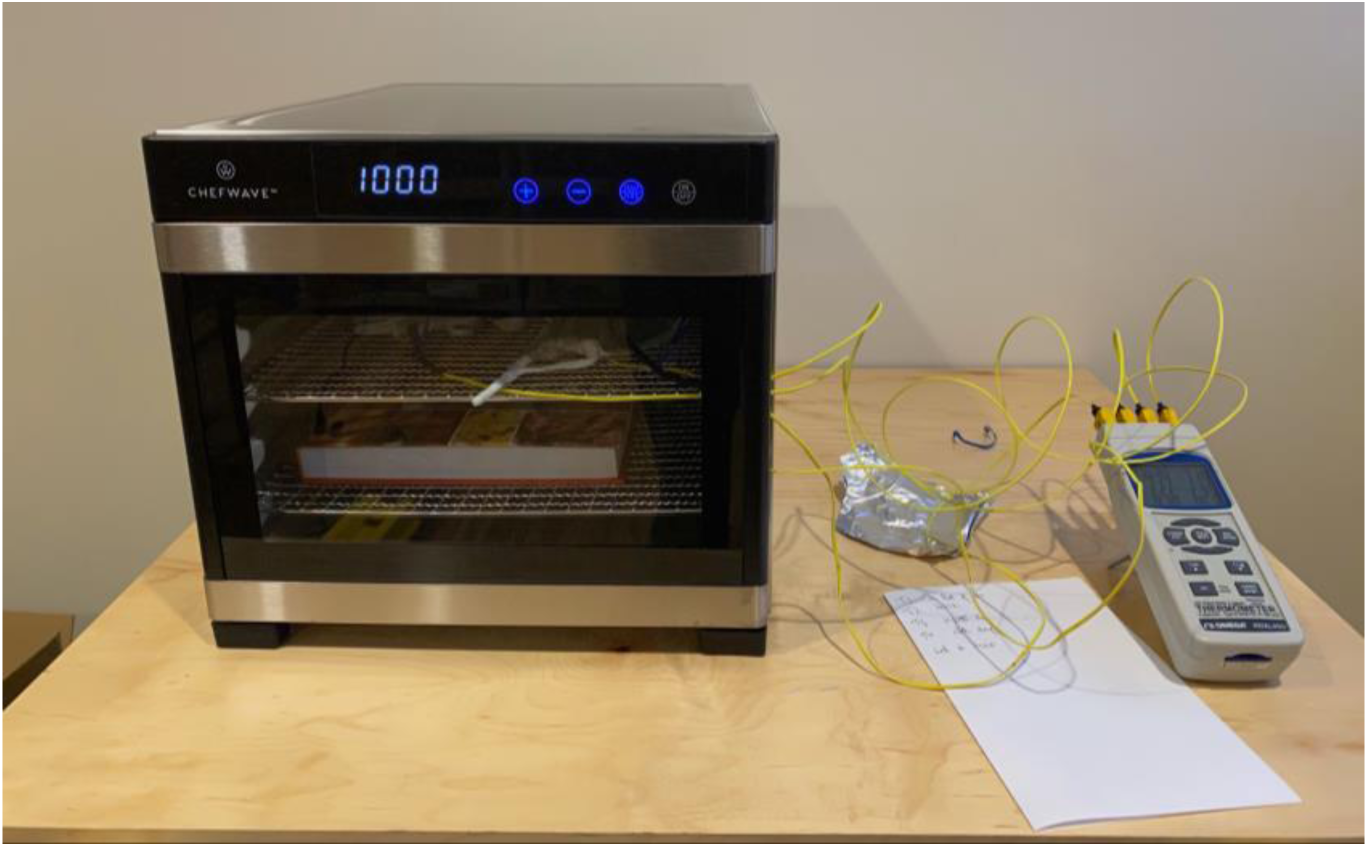
Food dehydrator

**Figure 4.**
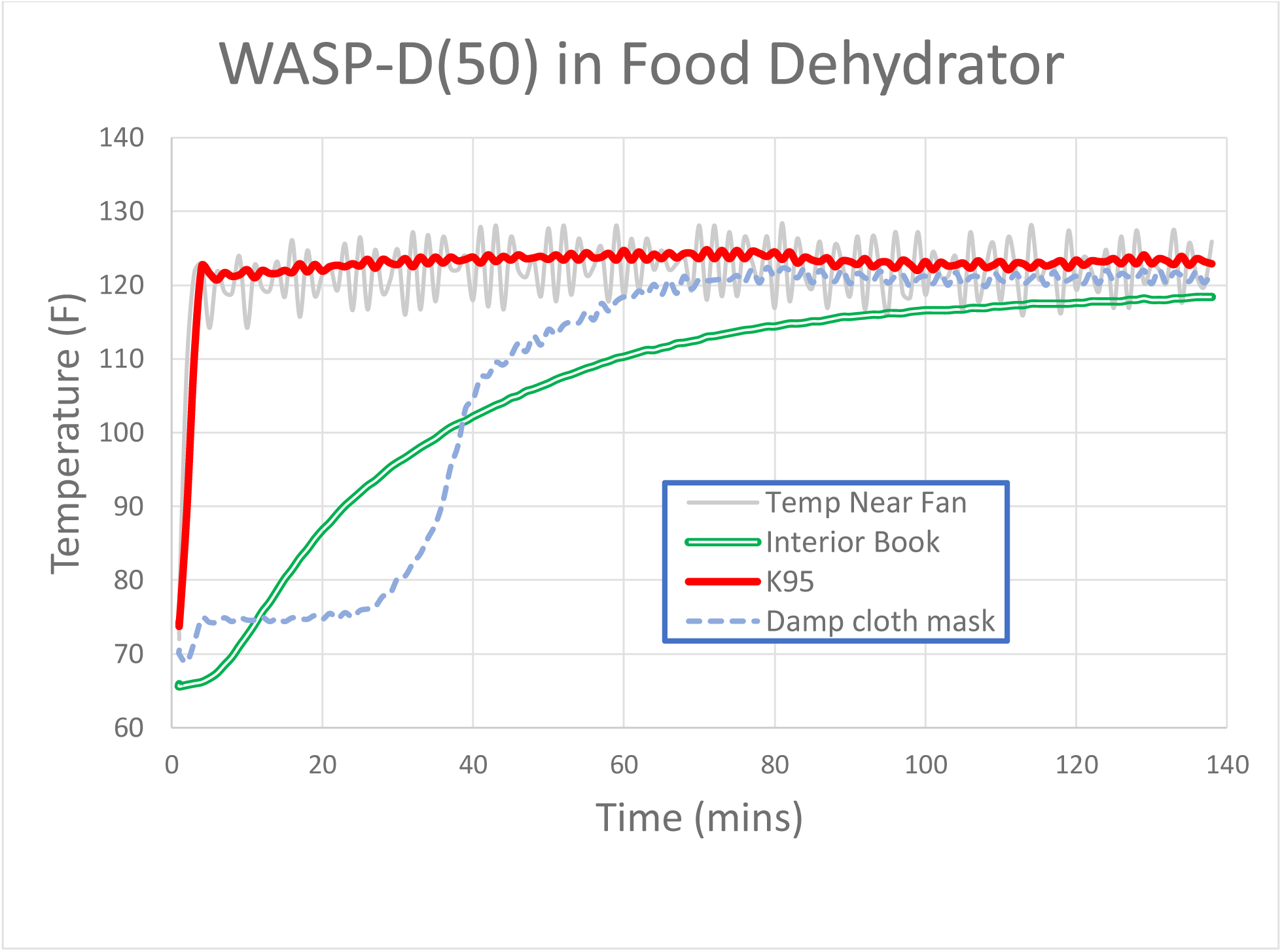
Time/Temperature curve inside dehydrator. Light gray curve is the air temperature inside dehydrator (cyclic variation is a consequence of the temperature controller). The green double line is the temperature inside a 1” thick book. The red line is the temperature inside a folded N95 mask. The blue dotted line is temperature history of a moist cloth mask-note it initially evaporatively cools below the chamber temperature, until most of the moisture has been driven off.

### Conclusions and recommendations

On BoCoV test samples and MS-2 e-coli vectors, WASP-D(46) is expected to result in a 10^6^ viral load reduction after 12 hours at 115°F=46°C and 22% relative humidity. We expect a similar effect on the human beta coronavirus SARS-CoV-2. While we have not yet field tested this methodology, WASP-D offers the prospect of saving lives around the globe by allowing for the relatively safe re-use of scarce PPE, and we hope others will confirm and extend our results and protocol in Appendix II.

While the intended audience for WASP-D are LMIN, it does not escape our notice that domestic hospitals and individuals would benefit from a convenient method to decontaminate PPE, or even entire rooms and buildings. New strains, such as alpha, are more contagious. Simple cloth masks are inadequate to stem this variant’s spread, but N95 masks are not yet available in sufficient quantities to be a viable substitute.

We can imagine colleges might employ this system to decontaminate student masks- or libraries, to sanitize circulating books and DVDs. We also emphasize, in this world of internet misinformation, **and in no uncertain terms**, that WASP-D cannot be used to decontaminate people. A steam spa, or blowing hot air up the nose, is ineffective and possibly dangerous. More subtly, exponential viral decay is exquisitely sensitive to time and temperature. **100**°**F and 8 hours is not a substitute for 115**°**F and 12 hours**. A home cooking oven set on WARM will overshoot the 125°F upper safe temperature range (due to radiation from the exposed heating elements) and possibly damage the PPE. Extra time must be allotted to account for the cooling effect of evaporating moisture, or low PPE thermal conductivity (see Appendix I).

Prudent care must be taken when applying WASP-D to real-world situations. A potential decontamination protocol is offered in Appendix II for discussion purposes-please contact the authors for up-to-date guidance.

## Appendix I: Initial temperature offset in WASP-D

WASP-D(50) implemented in a Chefwave CW-FD food dehydrator.

Temperatures were measured at four locations:

- In the air near the convection fan
- In the center of a 1” thick hardcover book
- In the center of a flat-folded K95 mask
- Between two layers of a damp cloth face mask

The dehydrator was set at 122F∼50C. Note how the temperature cycles on a regular basis as the controller seeks to maintain the target operating point. As illustrated, the controller holds the air temperature to a ±5F band.

The book, with pages made of cellulose (an excellent insulator), requires nearly 2 hours to enter the control band (while a stack of books might take ten hours or more). Thus, for a planned 12 hour WASP-D(50) ***library*** decontamination cycle, an additional two hours must be addended to the 12-hour WASP-D cycle for unstacked books.

The folded K95 mask, which is thin and dry, reaches 120F in under two minutes.

The damp cloth mask evaporatively cools, depressing its temperature to 75F for 30 minutes. Once nearly dry, its temperature rapidly increases, and by one hour is within the temperature cycling band. Thus, for damp materials of this kind and dimensions, an hour should be addended to the 12 hour WASP-D cycle.

## Appendix II: Proposed decontamination procedure

This specific proposed process has not been field tested, but is based on calculation and analogy with existing protocols.

### Summary

WASP-D (**W**arm **A**nd **S**low **P**athogen-**D**econtamination) enables the safe and simple decontamination of PPE from SARS-CoV-2 without damage or reduction of PPE effectiveness. Only heat, time and ambient humidity are required to lower SARS-CoV-2 viral load by six orders of magnitude. Heat also has the advantage of penetrating areas that cannot be reached by vapors or UV light, thus simplifying the decontamination process and reducing operating costs. WASP-D is well suited for low-resource environments and to relieve temporary PPE shortages.

To avoid damage to PPE materials, temperatures are limited to a “warm” range between 110F-122F (43C-50C), for a decontamination interval of 12hrs (at 46C/115F). Above 50C, or in the presence of UV-C, steam or chemical sanitizers, PPE elasticity or tear strength may be compromised, limiting reuse and possibly affecting its ability to exclude pathogens.

At these “warm” temperatures, some but not all pathogens will be diminished. Thus an “index’ system where decontaminated PPE are returned to their original wearer is prudent. Never the less, in situations where SARS-CoV-2 is the primary infectious agent, WASP-D can relieve PPE shortages and improve patient and health system outcomes.

### Caveats

The 12 hours cycle is predicated on thin, dry PPE. Wet PPE will evaporatively cool until it dries out. In a low humidity room this cooling can be substantial - perhaps 10C or more. And could take (in the case of a wrung-out sponge) five hours or more to dry, perhaps days for moist bedding. Similarly, if WASP-D is used to decontaminate thick, heavy insulating material, such as books or mattresses, extra time will be required. A single book might take two hours to reach 43C in a 46C room, while a stack of books (due to the insulation value of cellulosic paper) might take a day or more. These heating time delays must be added to the 12 hour WASP-D cycle. That is, if it takes 6 hours for a moist sponge to reach 43C, the entire WASP-D cycle is 18hrs=12hrs +6hrs.

### Facility Design

The WASP-D decontamination facility can take multiple forms, depending on local skills and resources. For example, a clean exterior shed. A greenhouse. A spare storage room. A shipping container. A car in the hot sun with closed windows and ventilation fan ON. Multiple WASP-D facilities allows for overlapping decontamination cycles.

Heat sources include sunlight (e.g. a greenhouse or black-painted shipping container), and electric heaters (ceramic heaters or oil radiators are preferred for fire safety and to prevent PPE exposure to infra-red radiation emanating from red-hot electric heating wires). Place heater inside facility near air inlet duct to pre-heat ventilation air.

Ideally, PPE should be stored on wire shelves (commercial, or locally built with chicken wire), with each shelf dedicated to PPE from one health worker. While gowns may be folded to save space, try to avoid contact between the presumably cleaner PPE interiors with the more highly exposed exterior surfaces. *Unlike UV-C and vaporized peroxide, heat will penetrate stacked masks and gowns, and in some cases, these denser arrangements of non-indexed PPE are expedient forms of decontamination*.

A remote thermometer to read the interior temperature is required--multiple thermometers, distributed within the facility, are preferred. **One thermometer probe should be placed inside the thickest (and/or moistest) PPE to confirm the decontamination temperature has been reached**. Temperatures should be maintained in the range of 110F to 125F (42C to 53C), preferably 115F (46C). Lower temperatures may not kill pathogens, and higher temperatures may damage some (but not all) PPE.

A ventilation fan to prevent mold and fungal growth is important. Preferably, this fan should blow air throughout the facility and exit via an exhaust vent or pipe which is oriented away from human contact or direct inhalation. One to two air exchanges an hour should be sufficient. Make sure heaters and wall insulation are adjusted to maintain target temperatures. In the absence of fans, natural convection from air registers near the floor of the facility to ceiling exhaust vents may be sufficient. However, in this case the entire facility must be located away from accidental human contact. For ventilation to be effective, do not over-stuff the facility with PPE.

### Protocol Steps

1. PPE exteriors are more likely highly contaminated than the interior. Thus, care should be taken to prevent individual PPE from touching each other. This step is not required if PPE is not indexed.
2. After PPE are stored in the WASP-D facility, the heat source is activated, and the ventilation fan triggered. Internal temperatures should be monitored during the 12 hour decontamination cycle to confirm it maintains 114F-120F (45C-49C), nominally 115F/46C.
3. Additional decontamination time may be required beyond 12 hours. Moist or sweaty PPE will evaporatively cool, and until it reaches ambient humidity, and may plateau 10C or more below ambient temperature. Thick materials gathered in a heap will self-insulate, delaying heat transfer to the center of the pile. ***WE HIGHLY RECOMMEND* WHEN APPLYING WASP-D ON THICK DAMP MATERIALS, SUCH AS MATTRESSES, TO INSERT A DIGITAL THERMOMETER PROBE INTO THE MIDDLE OF THE OBJECT AND CONFIRM IT HAS REACHED 114F (45C) BEFORE STARTING THE 12 HOUR DECONTAMINATION COUNTDOWN**
4. If for some reason the temperature drops (e.g. loss of heater power or sunlight) for a time interval, the WASP-D decontamination cycle should be increased by the same amount, plus time to return to target temperature. In-situ temperature measurement within the PPE are the best guide to confirm the PPE has reached target temperature of 115F (46C) or above. At which point the 12 hour cycle begins/continues.
5. A log of temperatures vs time during the pre-heat and 12 hour decontamination cycle should be maintained. Temperature swings ABOVE 50C (122F) should be avoided to limit thermal degradation of the PPE
6. After the WASP-D cycle completes, the facility may be opened. Decontaminated PPE may be collected by their indexed users directly from dedicated facility staff or stored in labeled and clean paper bags for later use.
7. As a precaution, a representative sample of PPE should be swabbed and tested to confirm pathogens levels are below target.
8. The facility should be thoroughly cleaned with soap/and or bleach and water and allowed to dry between cycles. This reduces cross-contamination from pathogens which are less heat-sensitive.

As every WASP-D system will be vary in configuration, we cannot guarantee this protocol is effective in all situations. Please implement carefully and use at your own risk. Please share your field experience with the authors so we can update and improve this advice.

## Footnotes

1 https://www.fda.gov/medical-devices/coronavirus-covid-19-and-medical-devices/medical-device-shortages-during-covid-19-public-health-emergency#shortage

2 https://www.cbsnews.com/news/face-mask-cdc-guidelines-covid-high-demand/

3 https://www.ebmt.org/low-middle-income-country-lmic-membership

4 https://www.cdc.gov/coronavirus/2019-ncov/hcp/ppe-strategy/decontamination-reuse-respirators.html

5 https://www.fda.gov/media/138284/download

6 https://www.battelle.org/inb/battelle-ccds-for-covid19-satellite-locations

7 https://en.wikipedia.org/wiki/Ozone_cracking

8 https://www.n95decon.org/publications#heat

9 https://www.cdc.gov/coronavirus/2019-ncov/hcp/non-us-settings/emergency-considerations-ppe.html

10 http://textbookofbacteriology.net/Listeria.html

11 http://textbookofbacteriology.net/staph.html

12 *Urban aerosols harbor diverse and dynamic bacterial populations*, Eoin L. Brodie, Todd Z. DeSantis, Jordan P. Moberg Parker, Ingrid X.Zubietta, Yvette M. Piceno, Gary L. Andersen, Proceedings of the National Academy of Sciences Jan 2007, 104 (1) 299-304; DOI:10.1073/pnas.0608255104

13 https://www.n95decon.org/implementation#time

14 https://www.fda.gov/media/138362/download

15 Stability of SARS-CoV-2 on Critical Personal Protective Equipment, Samantha B Kasloff, James E Strong, Duane Funk, Todd A Cutts, medRxiv 2020.06.11.20128884; doi:https://doi.org/10.1101/2020.06.11.20128884

16 The effect of temperature and humidity on the stability of SARS-CoV-2 and other enveloped viruses, Dylan H. Morris, Kwe Claude H. Yinda, Amandine Gamble, Fernando W. Rossine, Qishen Huang, Trenton Bushmaker, Robert J Fischer, M. Jeremiah Matson, Neeltje van Doremalen, Peter J Vikesland, Linsey C. Marr, Vincent Munster, James O Lloyd-Smith, bioRxiv 2020.10.16.341883; doi: https://doi.org/10.1101/2020.10.16.341883 This pre-print indicates less than a 45 minute half-life at 40C for a broad range of pathogens in a saliva matrix deposited on plastic test coupons.

17 https://www.tis-gdv.de/tis_e/containe/klima/klima-htm/; David Leinberger, Temperature & humidity in ocean containers. Technical report, Xerox Corporation, 2006.

18 Fischer RJ, Morris DH, van Doremalen N, et al. Effectiveness of N95 Respirator Decontamination and Reuse against SARS-CoV-2 Virus. *Emerging Infectious Diseases*. 2020;26(9):2253-2255. doi:10.3201/eid2609.201524

19 https://multimedia.3m.com/mws/media/1538979O/3m-disposable-respirator-1860-1860s-technical-data-sheet.pdf

20 https://genuineideas.com/ArticlesIndex/stallbbq.html

21 doi: 10.1021/acsnano.0c06565, DOI: 10.1128/AEM.02291-09, DOI: 10.1128/mSphere.00441-20

22 https://www.epa.gov/sites/production/files/2016-09/documents/emerging_viral_pathogen_program_guidance_final_8_19_16_001_0.pdf

23 ASTM International. *E2197-17e1 Standard Quantitative Disk Carrier Test Method for Determining Bactericidal, Virucidal, Fungicidal, Mycobactericidal, and Sporicidal Activities of Chemicals*. West Conshohocken, PA; ASTM International, 2017. doi: https://doi.org/10.1520/E2197-17E01

24 ASTM International. *E1052-20 Standard Practice to Assess the Activity of Microbicides against Viruses in Suspension*. West Conshohocken, PA; ASTM International, 2020. doi: https://doi.org/10.1520/E1052-20

25 https://doi.org/10.1002/9780471729259.mc15e02s37; Hasoksuz M., Vlasova A., Saif L.J. (2008) Detection of Group 2a Coronaviruses with Emphasis on Bovine and Wild Ruminant Strains. In: Cavanagh D. (eds) SARS- and Other Coronaviruses. Methods in Molecular Biology (Methods and Protocols), vol 454. Humana Press, Totowa, NJ. https://doi.org/10.1007/978-1-59745-181-9_5

26 Benfield DA and Saif LJ, J Clin Microbiol. 1990 Jun;28(6):1454-7. doi: 10.1128/JCM.28.6.1454-1457.1990. *Cell culture propagation of a coronavirus isolated from cows with winter dysentery*.

27 Lindenbach BD “Measuring HCV infectivity produced in cell culture and in vivo”, Methods Mol Biol. (2009) 510:329-36

24 Reference: Yang, L. et al. (2016). An improved plating assay for determination of phage titer. African Journal of Biotechnology. 15(23): 1078-1082.

